# Hot-spots and their contribution to the self-assembly of the viral capsid: *in-vitro* validation

**DOI:** 10.1101/724146

**Authors:** Alejandra Gabriela Valdez-Lara, Mariana Andrade-Medina, Josué Alejandro Alemán-Vilis, Aldo Adrián Pérez-Montoya, Nayely Pineda-Aguilar, Eduardo Martínez-Guerra, Abigail Roldán-Salgado, Paul Gaytán, Mauricio Carrillo-Tripp

**Affiliations:** Biomolecular Diversity Laboratory, Centro de Investigación y de Estudios Avanzados del Instituto Politécnico Nacional Unidad Monterrey, Vía del Conocimiento 201, Parque PIIT, C.P. 66600, Apodaca, Nuevo León, México; Centro de Investigación en Materiales Avanzados Unidad Monterrey, Parque PIIT, C.P. 66600, Apodaca, Nuevo León, México; Departamento de Ingeniería Celular y Biocatálisis, Instituto de Biotecnología, Universidad Nacional Autónoma de México, Av. Universidad 2001, Col. Chamilpa, Cuernavaca, Morelos 62210, México

**Keywords:** molecular biophysics, molecular biology, hot-spot prediction, CCMV, Thermal Shift Assay, TSA, Differential Scanning Micro-Calorimetry

## Abstract

The viral capsid is a macromolecular complex formed by self-assembled proteins (CPs) which, in many cases, are biopolymers with an identical amino acid sequence. Specific CP-CP interactions drive the capsid self-assembly process. However, it is believed that only a small set of protein-protein interface residues significantly contribute to the formation of the capsid; the so-called “hot-spots”. Here, we investigate the effect of *in-vitro* point-mutations on the *Bromoviridae* family structure-conserved interface residues of the icosahedral Cowpea Chlorotic Mottle Virus, previously hypothesized as hot-spots. We study the self-assembly of those mutated recombinant CPs for the formation of capsids by Thermal Shift Assay (TSA). We show that the TSA biophysical technique is a reliable way to characterize capsid assembly. Our results show that point-mutations on non-conserved interface residues produce capsids indistinguishable from the wild-type. In contrast, a single mutation on structure-conserved residues E176 or V189 prevents the formation of the capsid while maintaining the tertiary fold of the CP. Our findings provide experimental evidence of the *in-silico* conservation-based hot-spot prediction accuracy. As a whole, our methodology provides a framework that could aid in the rational development of molecules to inhibit virus formation, or advance capsid bioengineering to design for their stability, function and applications.

## 1. Introduction

Two essential molecular components constitute a virus particle, or virion. One is the genetic material in the form of a single or double-stranded RNA or DNA molecule which encodes all the genes necessary for infection and replication. Second, a sophisticated protein shell, known as the viral capsid, whose function is to transport and protect the genetic material from the harsh environment. The capsid is assembled from multiple chemically identical copies of one, or a few, capsid proteins (CPs) which pack the viral genome and relevant accessory proteins *in-vivo*. The CP-CP interactions are critical for the formation, stability, and integrity of the viral capsid. The extent of interactions and the type of resulting capsids are encoded in the CP as they yield uniform capsids in geometry and size. Hence, analysis of protein-protein interactions in viral capsids within and across members of virus families will shed light on the fundamental principles of the molecular mechanisms involved in the process of protein self-assembly.

Some viral CPs spontaneously form empty capsids *in-vitro* when expressed in heterologous systems and assembled in the right conditions [1–4]. Those empty capsids are known as virus-like particles (VLPs). In recent years, VLPs have gained attention because of their potential use in biotechnology and medicine. On the other hand, the inhibition of protein-protein interactions (PPI) is now a reality for an increasing number of cases [5]. This strategy has seen solid advances toward near-future therapeutic applications. Currently, one approach in this direction is the search, or rational design, of small molecules or peptides that inhibit specific PPIs. However, there are several problems involved in this process. One of them is that, as opposed to an enzyme’s active site, large interfaces stabilize PPIs. Nonetheless, it is reasonable to think that a small molecule interacting at the position of the so-called “hot-spots” will compete with the binding of the associated subunit without necessarily covering the whole surface of interaction. Hence, knowing the accurate location and physico-chemical nature of the hot-spots is key to develop inhibitory strategies that target PPIs.

According to the general hot-spot hypothesis, from all the residues forming the protein-protein interface, only a few of them contribute significantly to the formation of the protein complex. After the observation that, for a given virus family, there is a set of interface residues which are conserved in the CP amino acid sequence and at the same time in their 3D space location on the quaternary capsid structure [6], we hypothesized that those residues should have a critical role in the kinetics of capsid formation. We then proposed a structure-conservation-based methodology for the identification of such residues in icosahedral viral capsids. The details of the method, as well as its benefits in comparison to other hot-spot prediction strategies, are described in a companion article.

Application of the conservation-based method to the *Bromoviridae* family resulted in the identification of six conserved interface residues. A representative member of that family is the icosahedral Cowpea Chlorotic Mottle Virus (CCMV). The tertiary structure of the CP of the CCMV combines both *α*-helix and *β*-sheet secondary structures (Fig.1 a). A total of 180 identical copies of the CP self-assemble to form the capsid, creating a complex network of CP-CP interfaces arranged in an icosahedral geometry. Such an arrangement is characterized by distinct symmetry axes (Fig. 1b). An important geometrical feature is that all the oligomer interfaces related by the same symmetry axis are equivalent in the quaternary structure of the capsid. Two of the structure-conserved interface residues on the CP of the CCMV, P99 and F120, are located in the CP-CP interface related by a 3-fold or a 5-fold symmetry axis of the capsid. On the other hand, residues E176, R179, P188, and V189 are located in the CP-CP interface related by a 2-fold symmetry axis. It has been shown that the 2-fold related homodimers (e. g., A2-B1), are the first step in the kinetics of capsid assembly in the case of CCMV [7], hence the motivation to focus on this type of oligomer (Fig. 1c).

**Figure 1.**
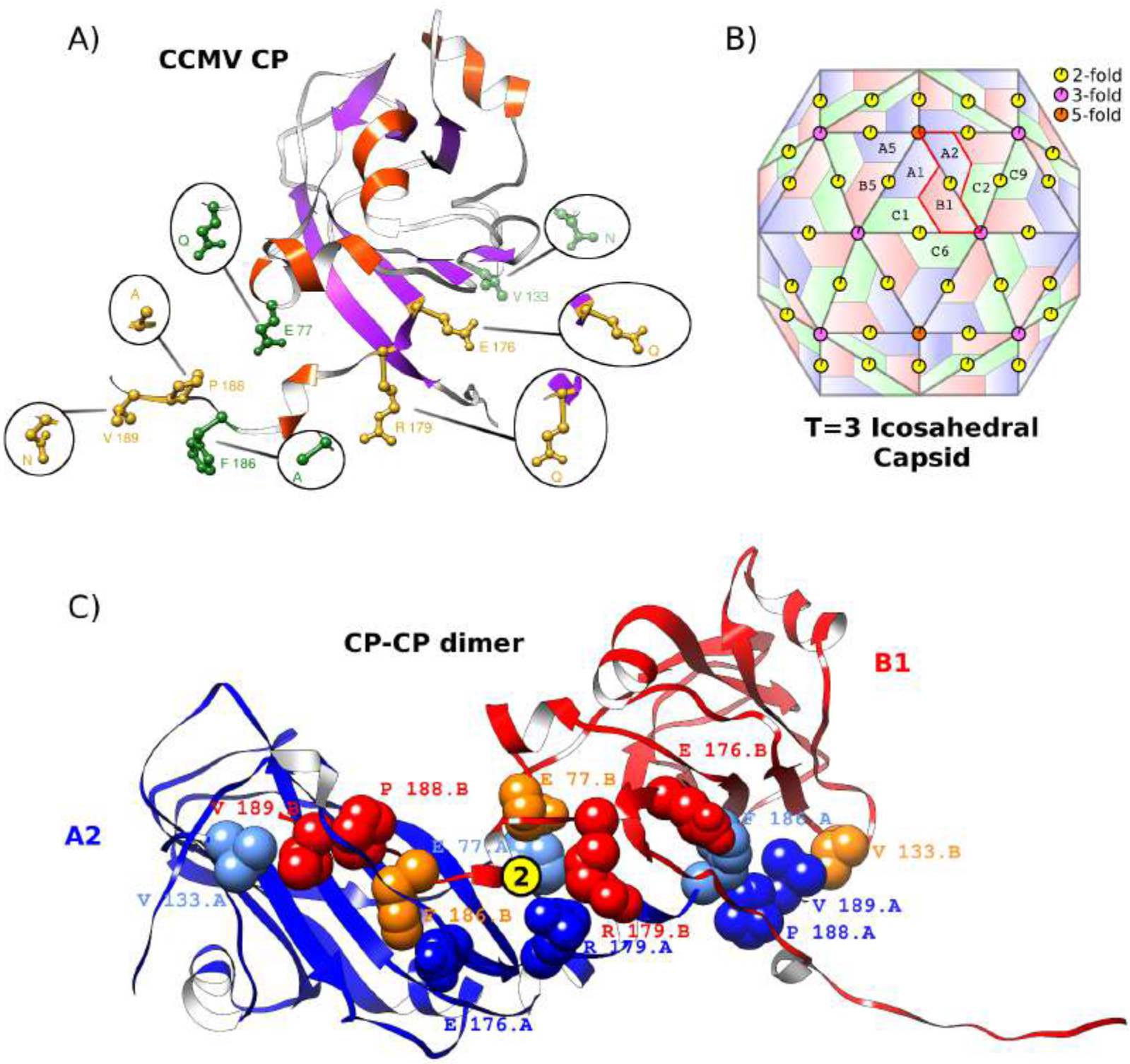
Molecular structure of the CP of the CCMV and capsid geometry. (**a**) Tertiary structure of the CP of the CCMV [14] (ribbon representation). Location of the 2-fold-related family structure-conserved interface residues (yellow side-chains) and three non-conserved interface residues (green side-chains). Corresponding point-mutations made are shown. The secondary structure is *α*-helix (orange) and *β*-sheet (purple). (**b**) 180 copies of the CP self-assemble to form the capsid. Here, each trapezoid represents one CP in the quaternary structure. Their color relates to local symmetry; a T=3 icosahedral geometry has distinct symmetry axes. Some of the 2-fold (yellow), 3-fold (magenta), and 5-fold (orange) axes are shown. A 2-fold-related homodimer is highlighted (A2-B1). All oligomers related by the same symmetry axis are *quasi*-equivalent in the quaternary structure (e. g., A2-B1 =A1-B5 = C1-C6 = C2-C9, etc.). (**c**) Quaternary structure of the 2-fold-related A2-B1 homodimer (blue-red ribbon representation), showing the location in the interface of the seven residues (VDW representation) corresponding to each CP; structure-conserved hot-spots (blue or red) and non-conserved interface residues (orange or cyan).

In this work, we conducted an *in-vitro* biochemical and biophysical study on the effect of individually mutating the 2-fold related structure-conserved interface residues of the CCMV. Regarding our experimental methodology, the Thermal Shift Assay (TSA) is a powerful tool that has been used to find protein stabilizing conditions [8,9], optimization of protein purification [10], study interacting ligands for drug discovery [11,12], and investigate biochemical properties of capsid proteins [13]. The setting up and running of TSA reactions is a simple process which allows for high-throughput studies. We implemented the TSA methodology on a qPCR machine to characterize the formation and stability of empty capsids when the recombinant CPs were in pH-dependent assembly conditions, as well as the stability of the CPs in monomeric state. We show systematically that the TSA accurately reproduces melting temperature values obtained with more expensive Differential Scanning Micro-Calorimetry experiments. Our results provide evidence that point-mutations on non-conserved interface residues do form empty capsids, indistinguishable from the wild-type. In contrast, point-mutations on two structure-conserved interface residues, E176Q or V189N, prevent the formation of the capsid while maintaining the CP tertiary fold. To the best of our knowledge, this is the first time an *in-vitro* validation is performed on theoretical hot-spot predictions of a viral capsid.

## 2. Results

We made *in-vitro* point-mutations to the CCMV CP in four structure-conserved interface residues (E176, R179, P188, and V189), two randomly-selected non-conserved interface residues (E77 and V133), and a non-conserved interface residue predicted to be a hot-spot by averaged energy-based approximation methods (F186) [15–17]. All of them are located at the interface formed around a 2-fold symmetry axis. The rationale followed in the point-mutations was to neutralize charges, change from non-polar to polar, or from big to small side chain, to disrupt all possible wild-type interactions. The final set was E176Q, R179Q, P188A, V189N, E77Q, V133N, and F186A. Figure 1 shows their location in the tertiary structure in one of the 180 CPs forming the capsid and in the quaternary structure of a 2-fold related dimer interface.

We produced the eight recombinant CP variants, wild-type and mutants, through an heterologous *E. coli* system in which we transformed the bacteria with plasmids with the corresponding modified CP genes. Previously reported CCMV CP expression and purification protocols were used [4]. TSA experiments allowed for the identification of the melting temperature of capsid disassembly, Tm_1_, and the melting temperature of CP denaturation, Tm_2_. For a given CP variant, the detection of both melting temperatures in the same sample through a TSA temperature scan gives quantitative evidence of the formation of capsids. We provide the software developed in this work to analyze the TSA data at https://github.com/tripplab/TSA-Tm. Details are given in the Methodology section.

### 2.1. Family structure-conserved interface residues E176 and V189 are critical for capsid formation

We conducted a series of TSA experiments to characterize the CCMV capsid self-assembly and the thermal stability of the point-mutated CPs. We studied two different conditions, namely, pH=4.8, in which wild-type CPs self-assemble into capsids, and pH=7.5, in which wild-type CPs stay at monomeric state. We found melting temperatures for capsid disassembly (Tm_1_) and CP denaturation (Tm_2_). Every assay included an independent measurement of Lysozyme as a positive control. A large number of experimental repetitions produced reliable and statistically significant results, summarized in Figure 2 (numerical values are reported in Table Sup. A1).

**Figure 2.**
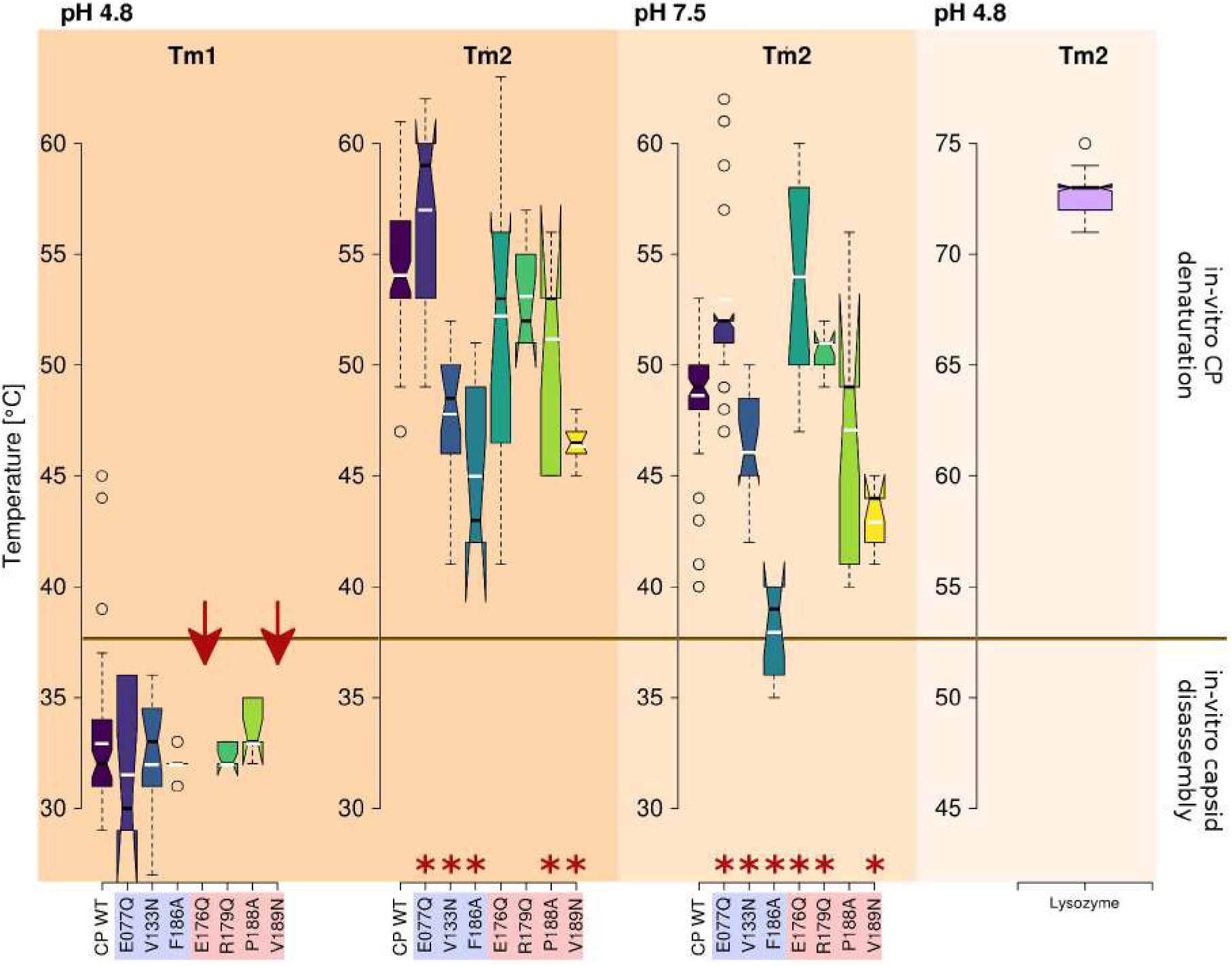
Self-assembly and thermal stability of CP variants of the CCMV. Statistical analysis (boxplots, black notch represents median, white bar average) of TSA experiments in two different conditions (pH=4.8 and pH=7.5) for the wild-type (WT), three non-conserved interface residues (blue box), and the four structure-conserved interface residues on the 2-fold-related CP-CP interface (red box). Melting temperatures were found for capsid disassembly (Tm_1_) and CP denaturation (Tm_2_) at pH=4.8. In the case of point-mutations E176Q and V189N, no capsids were formed (arrows). All TSA experiments included the Lysozyme independently at pH=4.8 as a positive control. * Statistical difference in the average values; *p-value* < 0.05 on a t-test with respect to WT.

Based on a consistent detection of Tm_1_ values, we found the formation of capsids at pH=4.8 in the case of WT CPs as expected [4]. The three point-mutations in the non-conserved interface residues, E77Q, V133N, and F186A, also form capsids. Two of the point-mutations in structure-conserved interface residues, R179Q and P188A, form capsids as well. None of those five mutants presented a Tm_1_ value significantly different from the characteristic capsid disassembly temperature of the wild-type CCMV (95% confidence intervals fall within the Tm_1_ value of the WT). Therefore, those five variants are indistinguishable from the wild-type, meaning that the quaternary structure and thermal stability of the capsid is not affected by those mutations. In contrast, structure-conserved point-mutations E176Q or V189N prevented the formation of the capsid, without disrupting the native fold of the CP.

As expected, none of the variants formed capsids at pH=7.5. To different degrees, all mutations seem to have affected the stability of the tertiary structure of the CP when in monomeric state, evidenced at high temperature values (Tm_2_). In the case of the non-conserved interface group, mutant E77Q increases, whereas mutants V133N and F186A decrease the Tm_2_ value with respect to WT. Family structure-conserved interface mutants E176Q and R179Q increase the Tm_2_ value, mutant P188A does not cause a detectable effect (higher experimental variability), and mutant V189N decreases the value.

Lowering the pH value from 7.5 to 4.8 increases the corresponding Tm_2_ value in all CP variants, as seen in the case of Lysozyme (Methods section). This observation is true in general, except for structure-conserved mutant E176Q, which has a large experimental variability. The pH value of 4.8 might be close to where the CP fold is the most stable (maximum in the Tm_2_ vs. pH profile). Nonetheless, structure-conserved point-mutations E176Q and R179Q do not affect the Tm_2_ at this pH value (95% confidence intervals fall within the Tm_2_ value of the wild-type).

## 3. Discussion

In this study, we provide experimental evidence for the validity and prediction power of the structural-conservation-based criteria for theoretical hot-spot identification. Our findings show that a single structure-conserved interface residue can play a significant role in the protein self-assembly mechanism, supporting the general hot-spot hypothesis [18]. Noteworthy, the structure-conserved residues we identified in the CCMV 2-fold related interfaces are significantly relevant only at the quaternary level as hypothesized, since the mutation to remove the physico-chemical contribution to the inter-molecular interactions of those residues do not destroy the tertiary protein fold but do inhibit capsid formation. This property does not hold for non-conserved interface residues, even when they have seemingly equivalent physico-chemical interactions, as mutations on such sites produce capsids indistinguishable from the wild-type. Hence, our *in-vitro* results are concomitant with the *in-silico* free energy study reported in the companion work.

By our structure-conserved definition of hot-spots, they are located in the inter-subunit interfaces and are conserved both in sequence and quaternary structure within all members of a virus family. In the case of the CCMV, member of the *Bromoviridae* family, the methodology identified six residues with such characteristics. Two of them are located in 3-fold and 5-fold related interfaces. The other four are located in the 2-fold related interfaces, and were studied here. Our hypothesis was that the point-mutation on any of them would either prevent the formation of capsids or significantly decrease the thermal stability of the particle. Here, we show that *in-vitro* mutations E176Q or V189N on the structure-conserved interface residues prevent the formation of the capsid, without unfolding the CP.

On the other hand, mutations R179Q and P188A on structure-conserved interface residues formed capsids with seemingly no distinction from the wild-type. Previous *in-silico* calculations estimated a similar contribution from those residues to the dimer binding free energy as that from structure-conserved residue V189 (45% of the binding free energy). When we compare the contacts involved in each one of the interface residues studied here (Table 1), we do not find any obvious feature or pattern that could explain why only V189N prevents capsid formation out of the three. For example, residue E176 has only one contact, with no apparent important interaction type. Nonetheless, binding free energy calculations showed that it is the structure-conserved interface residue which contributes the most to the protein complex (75% of the 2-fold related dimer binding free energy). Congruent with that calculation, *in-vitro* mutant E176Q does not form capsids. A similar mutation, but in a non-conserved site, E77Q, does not affect the quaternary structure and capsids are formed.

**Table 1.**
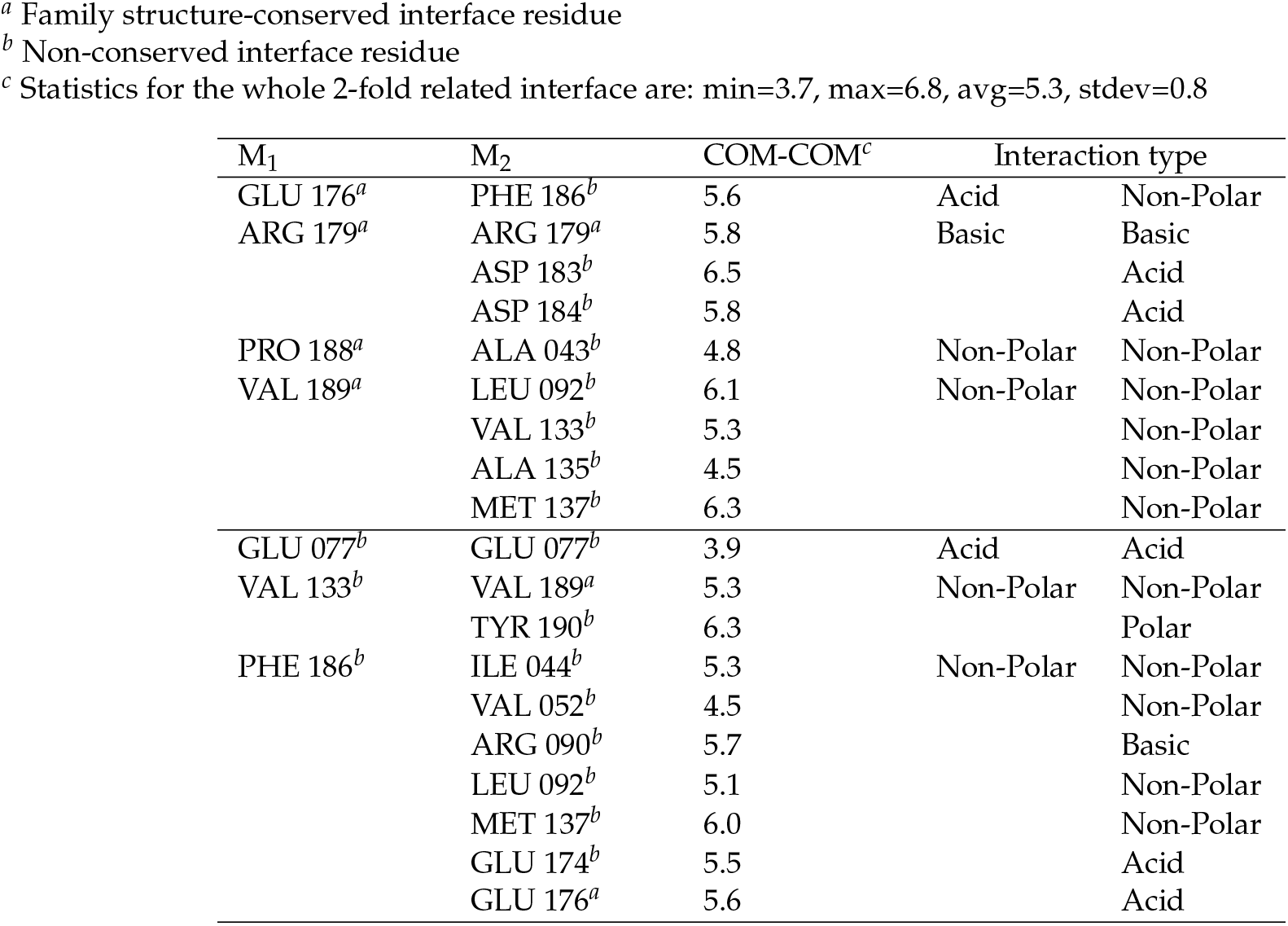
Contact table for the 2-fold related hot-spots and two non-conserved interface residues studied in this work. M_1_-M_2_ represents both A_2_-B_1_ and B_1_-A_2_ because of the homodimer symmetry. The distance between the center of mass (COM) between the residues in the pair is given in Å along with the physicochemical interaction type.

| M<sub>1</sub> | M<sub>2</sub> | COM-COM<sup>c</sup> | Interaction type |  |
| --- | --- | --- | --- | --- |
| GLU 176<sup>a</sup> | PHE 186<sup>b</sup> | 5.6 | Acid | Non-Polar |
| ARG 179<sup>a</sup> | ARG 179<sup>a</sup> | 5.8 | Basic | Basic |
|  | ASP 183<sup>b</sup> | 6.5 |  | Acid |
|  | ASP 184<sup>b</sup> | 5.8 |  | Acid |
| PRO 188<sup>a</sup> | ALA 043<sup>b</sup> | 4.8 | Non-Polar | Non-Polar |
| VAL 189<sup>a</sup> | LEU 092<sup>b</sup> | 6.1 | Non-Polar | Non-Polar |
|  | VAL 133<sup>b</sup> | 5.3 |  | Non-Polar |
|  | ALA 135<sup>b</sup> | 4.5 |  | Non-Polar |
|  | MET 137<sup>b</sup> | 6.3 |  | Non-Polar |
| GLU 077<sup>b</sup> | GLU 077<sup>b</sup> | 3.9 | Acid | Acid |
| VAL 133<sup>b</sup> | VAL 189<sup>a</sup> | 5.3 | Non-Polar | Non-Polar |
|  | TYR 190<sup>b</sup> | 6.3 |  | Polar |
| PHE 186<sup>b</sup> | ILE 044<sup>b</sup> | 5.3 | Non-Polar | Non-Polar |
|  | VAL 052<sup>b</sup> | 4.5 |  | Non-Polar |
|  | ARG 090<sup>b</sup> | 5.7 |  | Basic |
|  | LEU 092<sup>b</sup> | 5.1 |  | Non-Polar |
|  | MET 137<sup>b</sup> | 6.0 |  | Non-Polar |
|  | GLU 174<sup>b</sup> | 5.5 |  | Acid |
|  | GLU 176<sup>a</sup> | 5.6 |  | Acid |

Residue V189 is involved in four contacts inside an hydrophobic pocket. A change from a non-polar to a polar interaction will perturb the CP-CP interface. This effect seems to be significant only in the structure-conserved location. A similar mutation, but in a non-conserved site in a smaller pocket (two contacts), V133N, does not affect the capsid’s quaternary structure nor its thermal stability. Furthermore, the non-conserved interface residue F186, identified as a hot-spot by averaged energy-based approximation methodologies, fits into an even larger hydrophobic pocket (seven interface contacts). One would expect that a mutation on this site significantly perturbed the quaternary structure. Nonetheless, mutant F186A also produces capsids indistinguishable from the wild-type CP.

Point-mutation R179Q eliminates one repulsive and two attractive electrostatic interactions in the interface. Point-mutation P188A maintains a hydrophobic environment but increases the COM-COM distance of the interactions in this site, which should decrease the stability on the dimer. However, both mutations behave indistinguishably from the wild-type. We conclude that, if not relevant for the formation and stability of the capsid, there must be another, still unknown, evolutionary factor that has fixed them in sequence and quaternary structure.

In view of our findings, a similar study of the thermodynamic contribution of structure-conserved interface residues P99 and F120, which were excluded from this work due to being involved in protein interfaces other than the 2-fold related, is in progress. Most likely, their role will be in the formation of intermediate states in the process of capsid assembly, e. g., pentamer of dimers [19].

Finally, our results on the wide-range pH dependence of the denaturation temperature of Lysozyme by TSA showed an excellent agreement with DSC results. Such agreement is in line with previous reports that have shown a good correlation between the two techniques [20]. The fact that the TSA results are consistently lower than those from DSC was explained by the possibility that the dye used in TSA could be sensitive to temperature changes which locally affect the surface hydrophobicity and protein-solvent interactions, a factor that does not exist in DSC. Despite this, it is now accepted that the TSA method is reproducible, repeatable, and accurate; significant changes of a few degrees in Tm_*i*_ truly reflect a change on the investigated system.

## 4. Conclusions

We have shown in this and a companion work that the family-structure-conserved CCMV interface residues E176 and V189 largely contribute to the binding affinity and quaternary structure of the 2-fold related homodimer. This type of CP-CP complex is the first step in the capsid self-assembly process. Perturbations on those sites considerably affect the dimer and hence prevent capsid formation. In comparison to other hot-spot prediction strategies, our *in-vitro* mutagenesis results and thermal stability analysis prove that the conservation-based methodology is a more reliable strategy to identify residues with a significant contribution to the formation and stability of the protein complex.

## 5. Materials and Methods

### 5.1. CCMV recombinant CP variants are readily produced in an E. coli heterologous system

We corroborated correct in-frame cloning of all variants by sequencing the pET19b plasmid. Then, we ran Agarose gel electrophoresis resulting from subsequent PCR amplification of the CP wild-type and the six point-mutation genes using specific forward and reverse primers which amplified a 577 bp final product. After cell transformation, CCMV CP variants were individually expressed in *E. coli*. We verified the presence of the protein in each fraction of the purification process by monitoring its migration behavior in an SDS-PAGE gel at 12%. Most of the recombinant protein was retained after passing the bacterial clarified by the affinity column. The CP+His-TAG was eluted in 1.5 mL fractions after washing with small amounts of imidazole. Thus, other residual *E. coli* proteins that got trapped in the column were eluted. The CP wild-type (WT) has a molecular weight of ≈20 kDa [3] and a molecular extinction coefficient of 23,590 M^−1^ cm^−1^[21]. However, the CP+His-TAG variants migrate to a molecular weight of ≈24 kDa in the SDS-PAGE gel due to the inclusion of the histidine tag and all residues corresponding to the recognition site of the PreScission Protease^®^restriction enzyme. We have previously shown that the CP without the histidine tag migrates to the correct position. Furthermore, even with the histidine tag, CCMV VLPs are formed, i. e., the tag does not affect the self-assembly process [4]. The final concentration of the purified CP was ≈2 mg/mL (determined by Bradford protein assay) per 1 L of induced cell culture. We obtained all CCMV CP variants with high purity levels in most eluates, i.e., no other contaminating protein is detected. As a final verification, Western blotting with antibody against THE^*TM*^ His Tag Antibody was carried out for all variants. We summarize these results in Figure 3.

**Figure 3.**
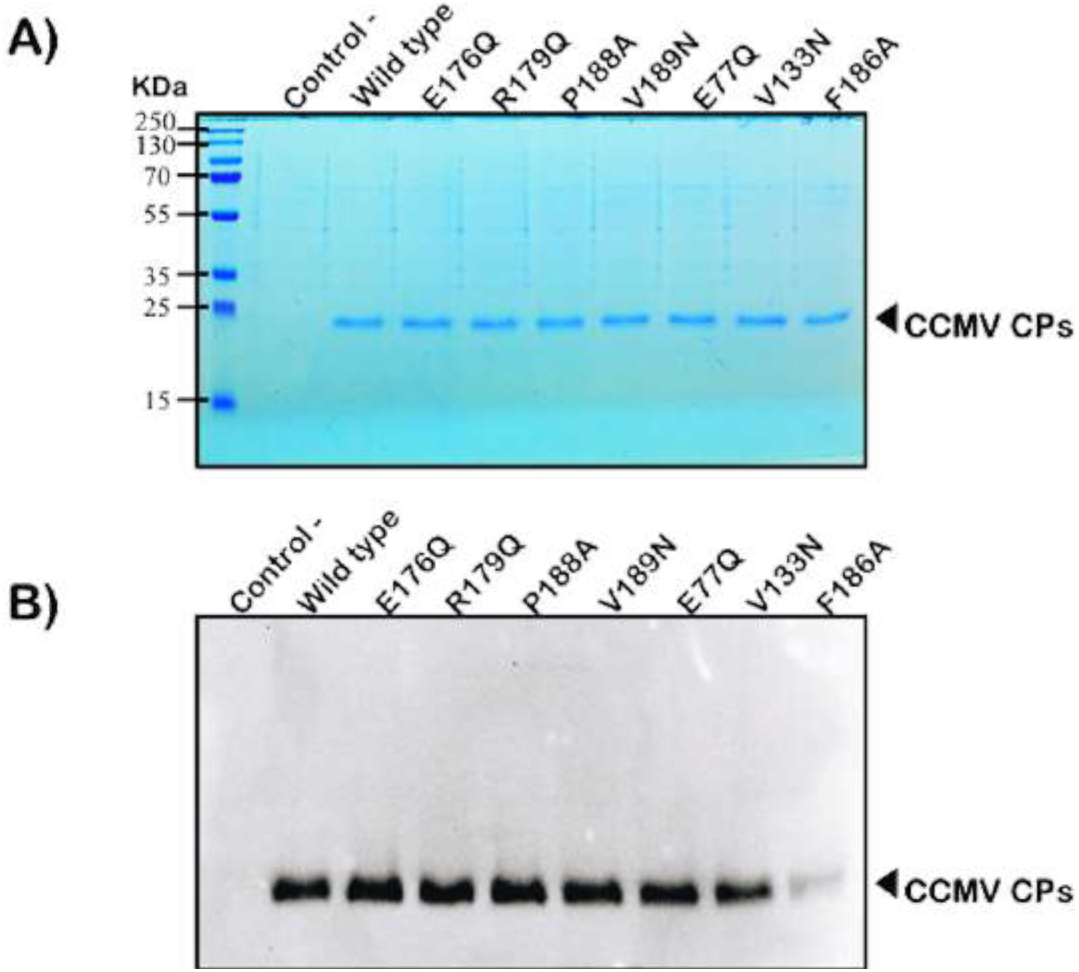
Recombinant CCMV CP variants. (**a**) Protein purification of the wild-type and the seven point-mutations. Molecular weight verified by SDS-PAGE electrophoresis analysis (visualized with Coomassie blue staining). (**b**) Recognition of the CPs his-tag by Western blotting with THE His Tag Antibody, mAb, Mouse (subtype IgG1). Non-transformed bacteria as a negative protein control is shown in both experiments.

#### 5.1.1. CCMV point-mutations *cp* genes construction

We cloned the nucleotide sequence of the native *cp* gene from CCMV into the pET19b plasmid [4]. Following the adapted PCR protocol previously reported, the amplification of the plasmid template containing the native *cp* gene was run in two separate PCR reaction. These reactions contained 1X High Fidelity buffer, 0.5 *μ*M dNTPs, 0.2 *μ*M of either forward or reverse primer, 10 ng/*μ*L of template plasmid, and 0.5 *μ*L of Phusion DNA polymerase (BioLabs ^®^) in a 25 *μ*L reaction. The PCR reaction was performed as follows: 1 cycle at 98 °C for 1:30 min, 10 cycles at 98 °C for 15 s, 60.4 °C for 30 s, 72 °C for 3:30 min, and 1 cycle at 72 °C for 5 min. After the first ten cycles, the two single-primer PCR products were combined in one test tube (giving a total volume of 50 *μ*L), and then the amplification was continued with 20 more cycles with the same conditions described above. The reaction was supplemented with 20 units of DpnI (BioLabs ^®^) and incubated overnight at 37 °C. The restriction enzyme DpnI digests the methylated template plasmid strands, leaving the newly synthesized strands intact. This product was purified and used to transform chemically competent *E. coli* DH5*α* cells. Plasmids were isolated from the resulting colonies and screened for the desired modification. We visualized PCR products and DNA plasmids by 1 % agarose (SIGMA Cat.#A9539) gel dyed with GelRed (BIOTIUM Cat.#41003). Finally, the positive clones were sequenced to confirm the desired modification and the absence of additional modifications.

#### 5.1.2. CCMV CP variants expression in *E. coli*

In order to express the CCMV CP variants, chemically competent cells of *E. coli*, Rosetta 2(DE3) pLysS ^®^(Novagen, Germany) were transformed with the CCMV point-mutated *cp* genes previously cloned into plasmid pET19b. This strain is resistant to chloramphenicol, and the plasmid pET19 confers resistance to ampicillin, carbenicillin, and related antibiotics. The transformed cells were grown in LB medium agar plate supplemented with carbenicillin (100 *μ*g/mL) and chloramphenicol (34 *μ*g/mL). One colony was selected, and a pre-inoculum was prepared in 100 mL of liquid LB medium supplemented with carbenicillin (100 *μ*g/mL) and chloramphenicol (34 *μ*g/mL). The culture was grown at 37 °C for 16 h and 200 rpm. This overnight culture was transferred to 1 L of fresh LB liquid medium, supplemented with carbenicillin and chloramphenicol, at the same concentration as before, and incubated at 37 °C and 200 rpm of agitation up to an optical density of 0.7 absorbance units. Finally, recombinant expression of the CCMV CP was induced with the addition of isopropyl-*β*-D-1-thiogalactopyranoside (IPTG), at a final concentration of 0.75 mM and incubation for 16 h at 25 °C and 200 rpm. Induced cells were collected by centrifugation at 3,000 × g at 4 °C for 30 min. The bacterial pellet was stored at −80 °C.

#### 5.1.3. CCMV CP variants purification by affinity chromatography

The pellet was re-suspended in 60 mL of lysis buffer (100 mM Tris-HCl, 500 mM NaCl, pH=6.0) supplemented with lysozyme (1 mg/mL final concentration) and phenylmethylsulfonyl fluoride (PMSF) to a final concentration of 1 mM, and incubated for 1 h at 4 °C with constant stirring. The solution was sonicated for 10 min, using 20 s pulses, at an intensity of 20% (QSONICA model Q500) with a standard 1/2 in diameter probe tip (#4220). Centrifugation at 13,500 x g was done for 30 min at 4 °C. The pellet was discarded, and the soluble fraction was filtered through a 0.45 *μ*m membrane and further loaded into a Pierce ^®^Cobalt affinity column (HisPur™ Cobalt Chromatography Cartridge) previously equilibrated with ten volumes of lysis buffer. Consecutive washes were made using three different buffers varying the imidazole concentration (100 mM Tris-HCl, 500 mM NaCl and L1 = 15 mM imidazole, pH=7.0; L2 = 20 mM imidazole pH=7.2; L3 = 35 mM imidazole, pH=7.3). Each wash was done with 20 volumes of the Cobalt affinity column. Recombinant CP was eluted using elution buffer (100 mM Tris-HCl, 500 mM NaCl, 500 mM imidazole, pH=8.0) in 2 mL fraction. Half of the recombinant protein was dialyzed for 16 h at 4 °C in 1 L of dialysis buffer (100 mM Tris-HCl, 500 mM NaCl, pH=7.5). To effectively retain the CP, a dialysis membrane of 14 kDa MWCO (MEMBRA-CEL ^®^MD44) was used. The concentration of the dialyzed protein was estimated using the Quick Start^®^Bradford Dye Reagent 1X (BIO-RAD Cat. #500-0205). The protein was stored at 4 °C before use.

#### 5.1.4. Protein separation and CCMV CP variants molecular weight signature

We verified the identity of the CCMV CP variants by molecular weight characterization. Sodium dodecyl sulfate (SDS) polyacrylamide gel electrophoresis assays were run for each of the different mutant CPs. Samples were mixed 4:1 with Laemmli 4x buffer (250 mM Tris-HCl pH=6.8, 8% SDS, 0.2% bromophenol blue, 40% glycerol and 20% 2-Mercaptoethanol). Total protein was denatured by heating at 97 °C for 5 min, cooled on ice for 5 min and loaded on the polyacrylamide gel at 12%. Each gel was run at 130 V for 1.5 h. Gels were stained using standard Coomassie Blue.

#### 5.1.5. Verification of the CCMV CP variants identity by Western Blot

Sodium dodecyl sulfate (SDS) polyacrylamide gel electrophoresis was performed as described before. Proteins were transferred to Immun-Blot ^®^PVDF membrane for protein Blotting (BIO-RAD Cat. #162 – 0177). Briefly, a “sandwich” was prepared with the following successive layers: i) three thicknesses of Mini Trans-Blot ^®^filter paper (BIO-RAD Cat. #1703932) 7.5 x 10 cm; ii) the SDS-polyacrylamide slab gel (with the stacking gel removed); iii) Immun-Blot ^®^PVDF membrane; iv) three more thicknesses of filter paper. All components in contact with the slab gel were previously wet in the transfer solution (10 mM Tris-base, 50 mM Glycine, 10% methanol). The blot was run using a Semidry Electroblotter from Thermo Fisher Scientific (Model HEP-1) at 15 V for 1 h using a BIO-RAD PowerPac ^®^HC power supply. Immediately after the transfer, the PVDF membrane was immersed in a wash buffer solution (100 mM Tris-HCl, 0.9% NaCl, 0.1% Tween20 pH=7.5) and incubated at 25 °C for 15 min, then it was briefly rinsed with two changes of wash buffer. The membrane was blocked for non-specific binding sites by immersion in Block buffer (100 mM Tris-HCl, 0.9% NaCl, 3% non-fat dry milk, pH=7.5) at 25 °C for 1 h. The recombinant protein was detected using the primary antibody THE™ His Tag Antibody, mAb, Mouse (subtype IgG1) GenScript (Cat. #A00186) diluted 1:3000 in buffer (100 mM Tris-HCl, 0.9% NaCl, 0.05% Tween20, 0.5% non-fat dry milk, pH=7.5). The membrane was incubated at 4 °C for 16 h, washed and incubated again with HRP labeled secondary antibody diluted at the same dilution factor and buffer as the primary antibody. The detection was made using SuperSignal^®^West Dura Extended Duration Substrate (Thermo Scientific Cat. #34075) according to the manufacturer’s instructions. The blot was photographed 5 min after the substrate was applied using the Bio-Imaging System (MicroChemi Cat. #70 – 25 – 00).

### 5.2. CCMV capsids spontaneously self-assemble in pH=4.8

The correct formation of wild-type CCMV capsids was confirmed by tungsten negative-stain on a Scanning Electron Microscope (SEM). After expression and purification, we placed the protein sample in assembly buffer (pH=4.8) on a copper grid with a formvar/carbon-coated mesh. We found a high density of capsids having the characteristic ≈30 nm diameter (Figure 4).

**Figure 4.**
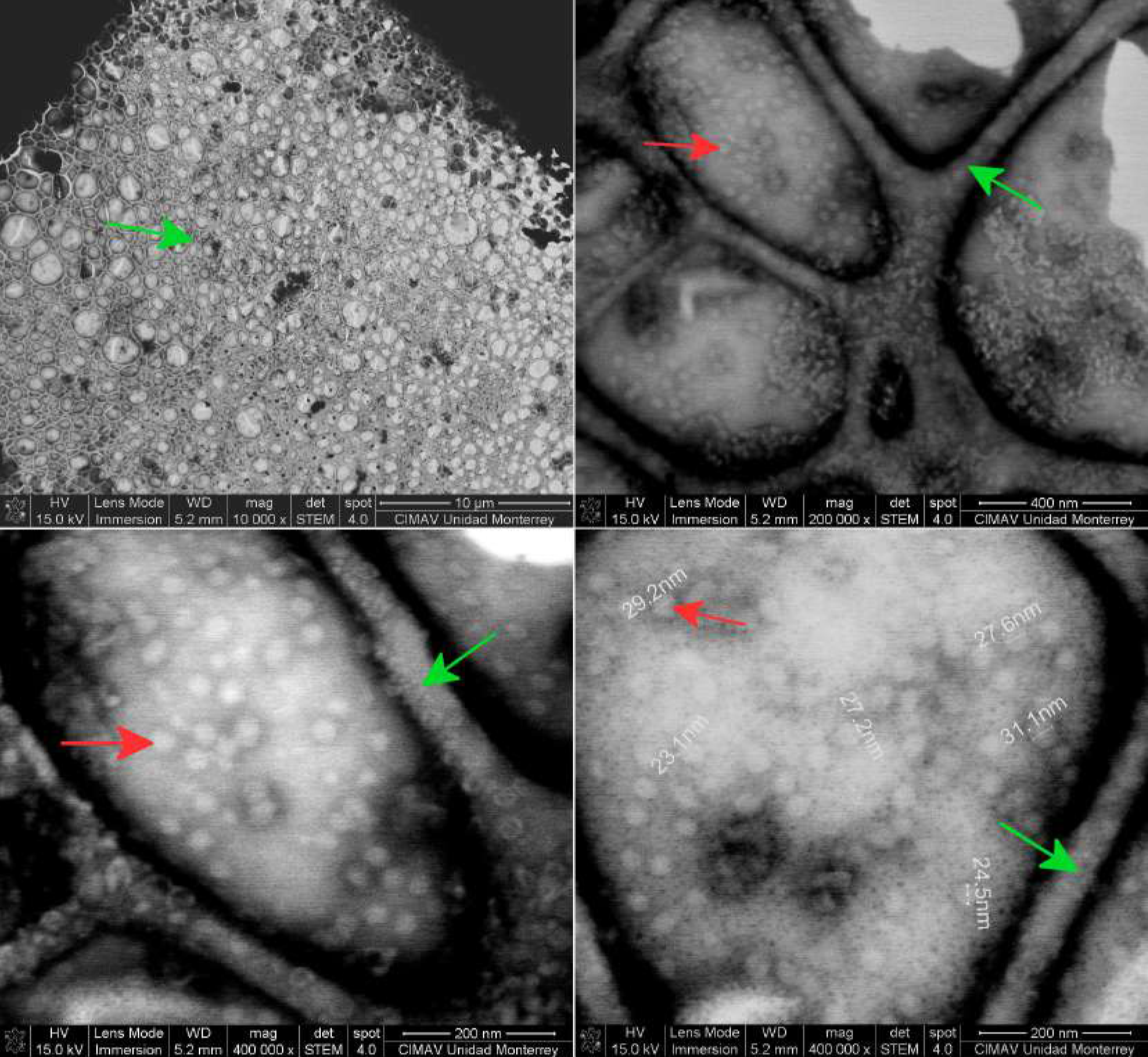
Virus-like particles of wild-type CCMV saw on a scanning electron microscope (SEM) by negative-stain. The sample was placed on a 300 copper grid with a formvar/carbon-coated mesh (green arrows). A high density of VLPs with a characteristic diameter of ≈30 nm are visible at magnifications of 200,000x and 400,000x (red arrows).

#### 5.2.1. *in-vitro* assembly of CCMV capsid variants

The self-assembly process of the different CP variants was induced in aliquots of purified protein placed separately in a dialysis membrane of 14 kDa MWCO (MEMBRA-CEL^®^MD44) and dialyzed simultaneously at 4 °C for 16 h against 1 L of assembly buffer (0.1 M Sodium acetate buffer, 500 mM NaCl, pH=4.8).

#### 5.2.2. Electron Microscopy

We confirmed capsid formation by direct observation in a Scanning Electron Microscope (SEM). Samples were taken after the assembly reaction and diluted to 0.15mg/mL. 3 *μL* of the diluted sample was placed on hydrophilic formvar coated mesh copper grids and incubated for 10min at room temperature to allow for capsid adherence. Excess sample was removed by absorption with filter paper. 3 *μ*L of phosphotungstic acid (PTA) at 2% was added on the formvar grid for negative staining. The sample was incubated for 10 min, and excess PTA was removed with filter paper. Grids were examined in a Scanning Electron Microscope Field Emission Model 200 Nova NanoSEM FEI Company.

### 5.3. Thermal characterization of CCMV CP variants self-assembly

The use of dyes like SYPRO Orange to monitor thermal denaturation of proteins by fluorescence detection provides a fast and inexpensive way for determination of protein stability using qPCR machines [22]. This technique is commonly referred to as Thermal Shift Assay (TSA). Here, we use such a methodology to study the CP variants self-assembly and capsid stability in solution. The capsid-dye sample is gradually heated (*T*) while measuring the fluorescence signal (*F*). The methodology enables for the detection of two characteristic melting temperatures; Tm_1_ corresponds to the capsid disassembly, while Tm_2_ corresponds to the protein thermal unfolding (denaturation). The specific value of Tm_*i*_ is defined as *max_i_*(*dF*/*dT*).

We developed a software, TSA-Tm, to analyze the raw data produced by TSA experiments. If the analyzed protein does not form oligomers of any kind, we will only find one distinctive peak on the first derivative of the fluorescence vs. temperature profile. On the other hand, when a protein self-assembles into oligomers, capsids in this case, we will find a peak corresponding to the protein complex disassembly, Tm_1_, and a second peak corresponding to the thermal denaturation of the protein, Tm_2_. We show an illustration of the second case in Figure Sup. A1.

In order to characterize the reliability of the TSA methodology, we compared it with previously reported results by Differential Scanning Micro-calorimetry (DSC) of Lysozyme [23]. We conducted a systematic analysis in a pH range of 1 to 12 and measured the Lysozyme denaturation temperature Tm_2_ behavior. We describe the specific conditions used for each pH value in Table Sup. A2. We fitted a third-order orthogonal polynomial model to the TSA data. The coefficients found were *Y* = 34.1464646 + 20.75628*X* − 3.17248*X*^2^ + 0.13604*X*^3^, with an R^2^=0.9251 and all the coefficient terms being statistically significant with a *p-value* < 0.001. Both DSC and TSA Tm_2_ values for Lysozyme fall within a 95% confidence interval of the model, demonstrating that both methods can detect the same quantitative trends in protein thermostability (Figure 5).

**Figure 5.**
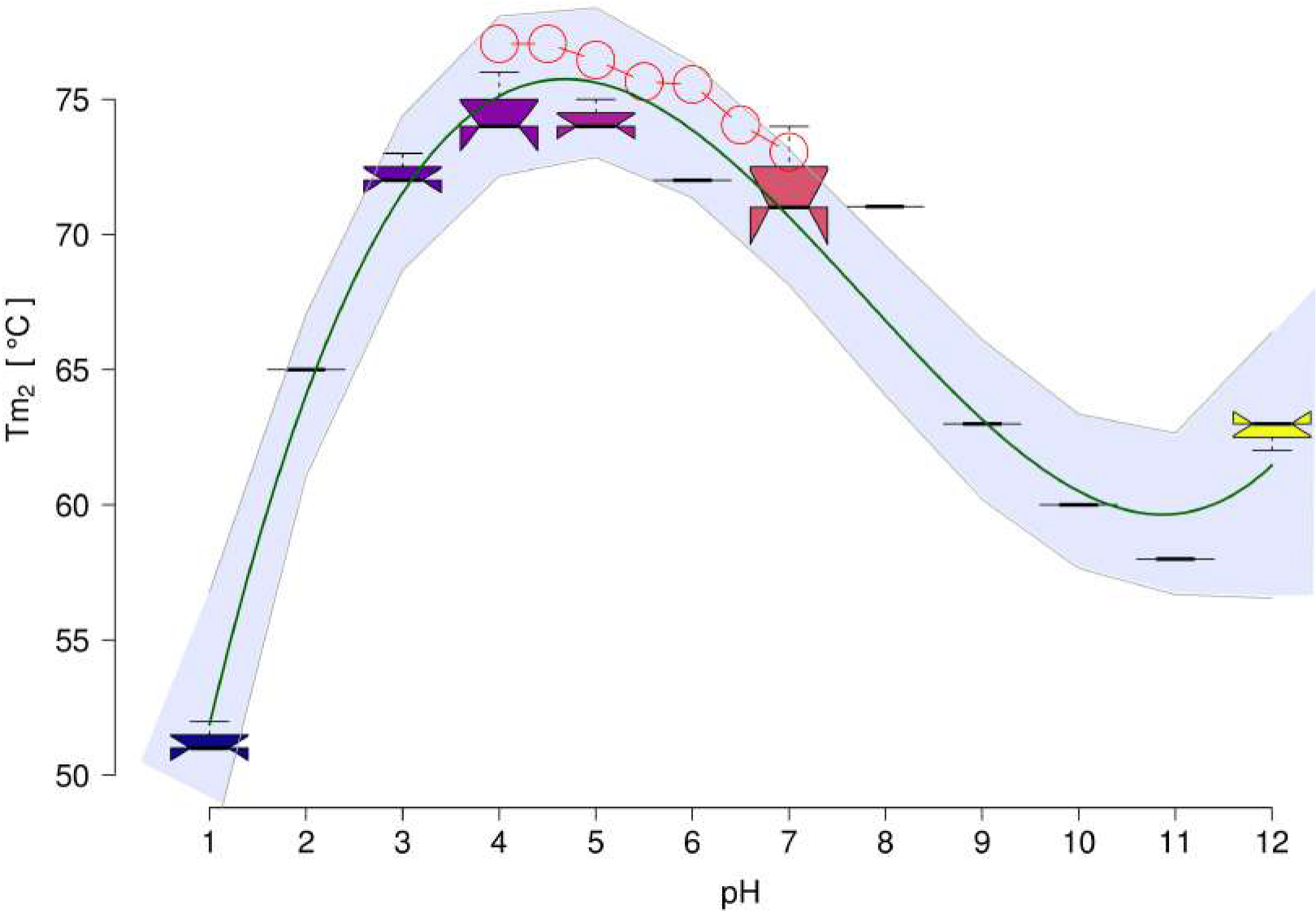
Denaturation temperature, Tm_2_, of Lysozyme as a function of pH value (boxplots). Previously reported values measured by Differential Scanning Micro-calorimetry (DSC) are shown in red circles [23]. A third-order orthogonal polynomial model was fitted to the TSA data (green line; R^2^=0.9251 with all the coefficient terms being statistically significant with a *p-value* < 0.001). The shaded region indicates 95% confidence intervals.

The protocol for a TSA experiment involves the incubation of CCMV CP variants with SYPRO Orange dye in PCR strip tubes. Each reaction contains the following components in a final volume of 30 *μ*L: 1x assembly buffer (0.1 M Sodium acetate buffer, 500 mM NaCl, pH=4.8), or 1x disassembly buffer (100 mM Tris-HCl, 500 mM NaCl, pH=7.5), 7.5 *μ*M of purified CCMV CP variants, and 6x SYPRO Orange. The TSA experiments were performed using a StepOne™ Real-Time PCR System (Applied Biosystems Cat. #4376357), with a temperature ramp starting at 25 °C and ending at 95 °C in 1 °C/min increments. The fluorescence signal was read at excitation and emission *λ* wavelengths of 490 nm and 585 nm respectively. An independent sample of Lysozyme (Sigma 62970) in a pH=4.8 buffer was included in each experiment run as a positive control. Raw data sets were exported from the qPCR machine for further analysis using the software we describe below.

### 5.4. Data analysis

The raw data produced by the readings of the qPCR machine are the fluorescence intensity as a function of cycles or temperature per well. Depending on the plate size and the number of optical channels, 48, 96, or more experiments per run can be made with information from several wavelength readings in a high-throughput fashion. Usually, the qPCR machine control software can export this raw data set. However, further analysis to obtain the melting temperature (Tm) can be a cumbersome process. We developed a Matlab program, TSA-Tm, to calculate such values. In addition to facilitate the analysis by reading and ordering the exported qPCR data in the XLSX formatted file, it provides a graphical interface where the user can choose the optical channel and set of wells to analyze, calculate the average and standard deviation of the fluorescence vs. temperature profile if more than one repetition was included in the experimental design, and numerically calculate *dF*/*dT*.The TSA-Tm open-source code and instructions to install and use the software can be obtained at https://github.com/tripplab/TSA-Tm. Once we obtained the Tm_*i*_ values for all the CP variants and conditions, we used the R software environment version 3.6.1 for statistical data analysis and plotting with RStudio version 1.1.463. The analysis R source code is described in Supplementary Information Section A.2.

## Author Contributions

Conceptualization, M.C.-T.; methodology, M.A.-M and M.C.-T.; software, A.A.P.-M.; validation, A.G.V.-L., J.A.A.-V. and M.A.-M; formal analysis, M.C.-T.; investigation, A.G.V.-L., J.A.A.-V., M.A.-M, N.P.-A., E.M.-G., A.R.-S., P.G. and M.C.-T.; resources, N.P.-A., E.M.-G.. P.G. and M.C.-T.; data curation, M.C.-T.; writing–original draft preparation, M.C.-T.; writing–review and editing, M.C.-T.; visualization, A.G.V.-L., A.A.P.-M., M.A.-M, N.P.-A. and M.C.-T.; supervision, M.A.-M and M.C.-T.; project administration, M.A.-M and M.C.-T.; funding acquisition, M.C.-T.

## Acknowledgments

The authors thank Dr. Bruno A. Escalante Acosta from CINVESTAV Unidad Monterrey for kindly providing access to the qPCR machine used in this study.

## Abbreviations

The following abbreviations are used in this manuscript:

CCMV: Cowpea Chlorotic Mottle Virus
CP: capsid protein
VLP: virus-like particle
PPI: protein-protein interactions
WT: wild-type
TSA: Thermal Shift Assay
DSC: Differential Scanning Micro-calorimetry
COM: center-of-mass

## Appendix A Supplementary data

### Appendix A.1 Supplementary figures and tables

**Figure A1.**
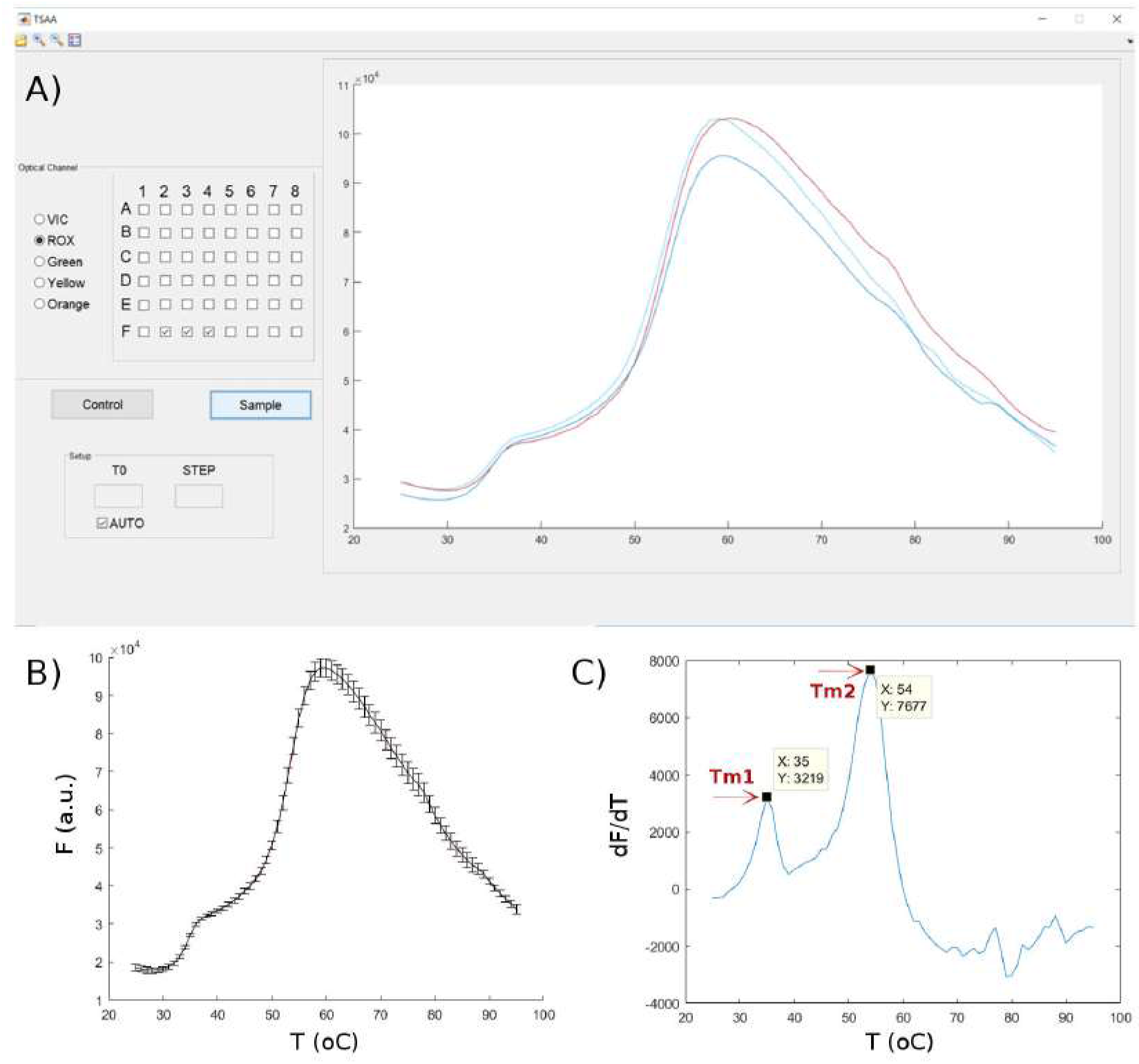
TSA-Tm analysis tool. A) The user can select the optical channel and the set of wells to analyze and visualize the results through a graphical interface. Then, B) subtraction of the blank (control) and averaging of repetitions, as well as C) calculation of *dF*/*dT* is automatic. The melting temperature values, Tm_*i*_, correspond to the peaks found in the first derivative of the fluorescence vs. temperature profile.

**Table A1.**
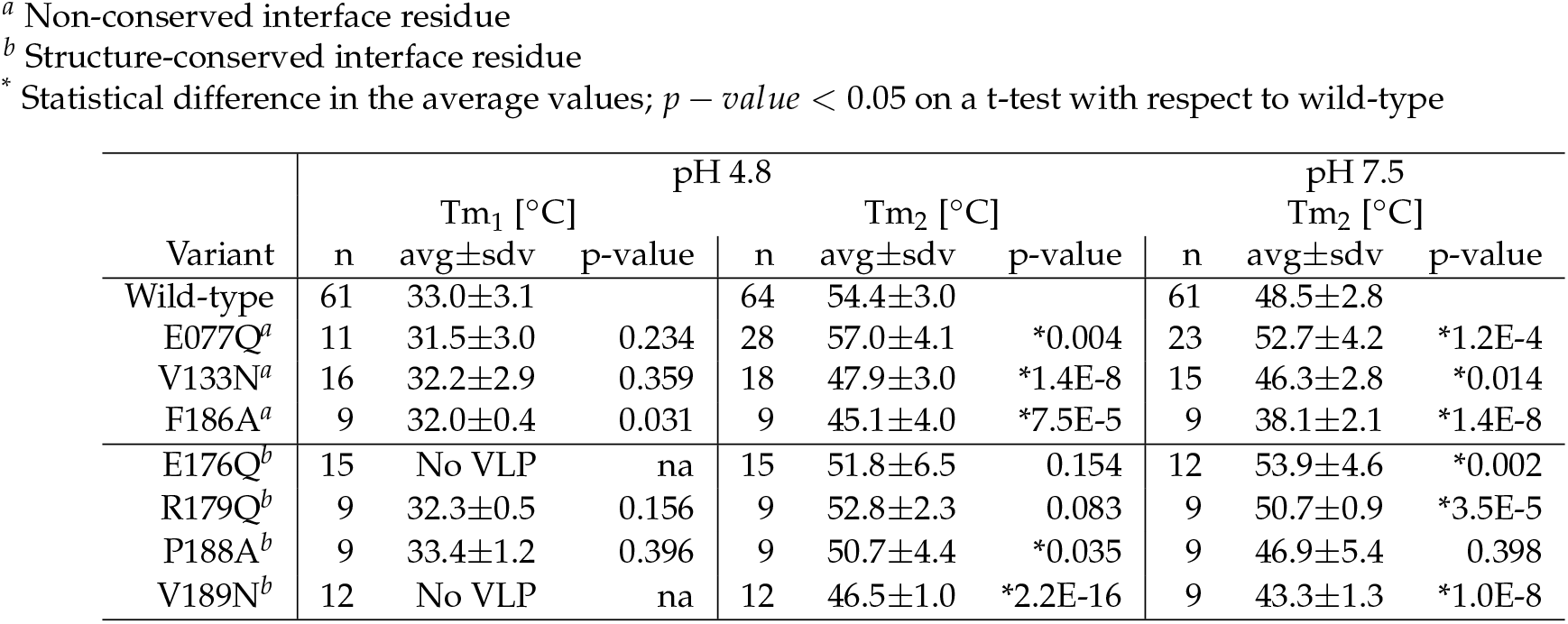
Thermal Shift Assay melting temperature (Tm_*i*_) values. Two different pH values were tested; 4.8 capsid self-assembly conditions, and 7.5 capsid disassembly conditions. Tm_1_ is the capsid melting temperature, and Tm_2_ is the CP unfolding temperature.

| Variant | pH 4.8 |  |  |  |  |  | pH 7.5 |  |  |
| --- | --- | --- | --- | --- | --- | --- | --- | --- | --- |
|  | n | Tm<sub>1</sub> [°C]<br>avg±sdv | p-value | n | Tm<sub>2</sub> [°C]<br>avg±sdv | p-value | n | Tm<sub>2</sub> [°C]<br>avg±sdv | p-value |
| Wild-type | 61 | 33.0±3.1 |  | 64 | 54.4±3.0 |  | 61 | 48.5±2.8 |  |
| E077Q<sup>a</sup> | 11 | 31.5±3.0 | 0.234 | 28 | 57.0±4.1 | *0.004 | 23 | 52.7±4.2 | *1.2E-4 |
| V133N<sup>a</sup> | 16 | 32.2±2.9 | 0.359 | 18 | 47.9±3.0 | *1.4E-8 | 15 | 46.3±2.8 | *0.014 |
| F186A<sup>a</sup> | 9 | 32.0±0.4 | 0.031 | 9 | 45.1±4.0 | *7.5E-5 | 9 | 38.1±2.1 | *1.4E-8 |
| E176Q<sup>b</sup> | 15 | No VLP | na | 15 | 51.8±6.5 | 0.154 | 12 | 53.9±4.6 | *0.002 |
| R179Q<sup>b</sup> | 9 | 32.3±0.5 | 0.156 | 9 | 52.8±2.3 | 0.083 | 9 | 50.7±0.9 | *3.5E-5 |
| P188A<sup>b</sup> | 9 | 33.4±1.2 | 0.396 | 9 | 50.7±4.4 | *0.035 | 9 | 46.9±5.4 | 0.398 |
| V189N<sup>b</sup> | 12 | No VLP | na | 12 | 46.5±1.0 | *2.2E-16 | 9 | 43.3±1.3 | *1.0E-8 |

**Table A2.** Buffer preparation to measure protein melting temperatures by the thermal shift assay method in a wide range of pH values.

| pH | buffer | preparation * |
| --- | --- | --- |
| 1 | KCl/HCl | 0.93g/0.77mL |
| 2 | KCl/HCl | 0.186g/0.043mL |
| 3 | Gly/HCl | 0.18g/0.048mL |
| 4 | AcONa/AcOH | 0.225g/0.54mL |
| 5 | AcONa/AcOH | 0.7g/ 0.21mL |
| 6 | Na<sub>2</sub>HPO<sub>4</sub>/NaH<sub>2</sub>PO<sub>4</sub> | 0.46g/1.48g |
| 7 | Na<sub>2</sub>HPO<sub>4</sub>/NaH<sub>2</sub>PO<sub>4</sub> | 1.94g/0.73g |
| 8 | TrizmaHCl/TrizmaBase | 1.081g/0.693g |
| 9 | NaCO<sub>3</sub>/NaHCO<sub>3</sub> | 0.115g/0.917g |
| 10 | NaCO<sub>3</sub>/NaHCO<sub>3</sub> | 0.143g/0.097g |
| 11 | NaHCO<sub>3</sub>[0.05M]/NaOH[0.1M] | 16.6mL/7.56mL |
| 12 | KCl[0.2M]/NaOH[0.2M] | 20mL/4.8mL |

### Appendix A.2 Analysis R source code

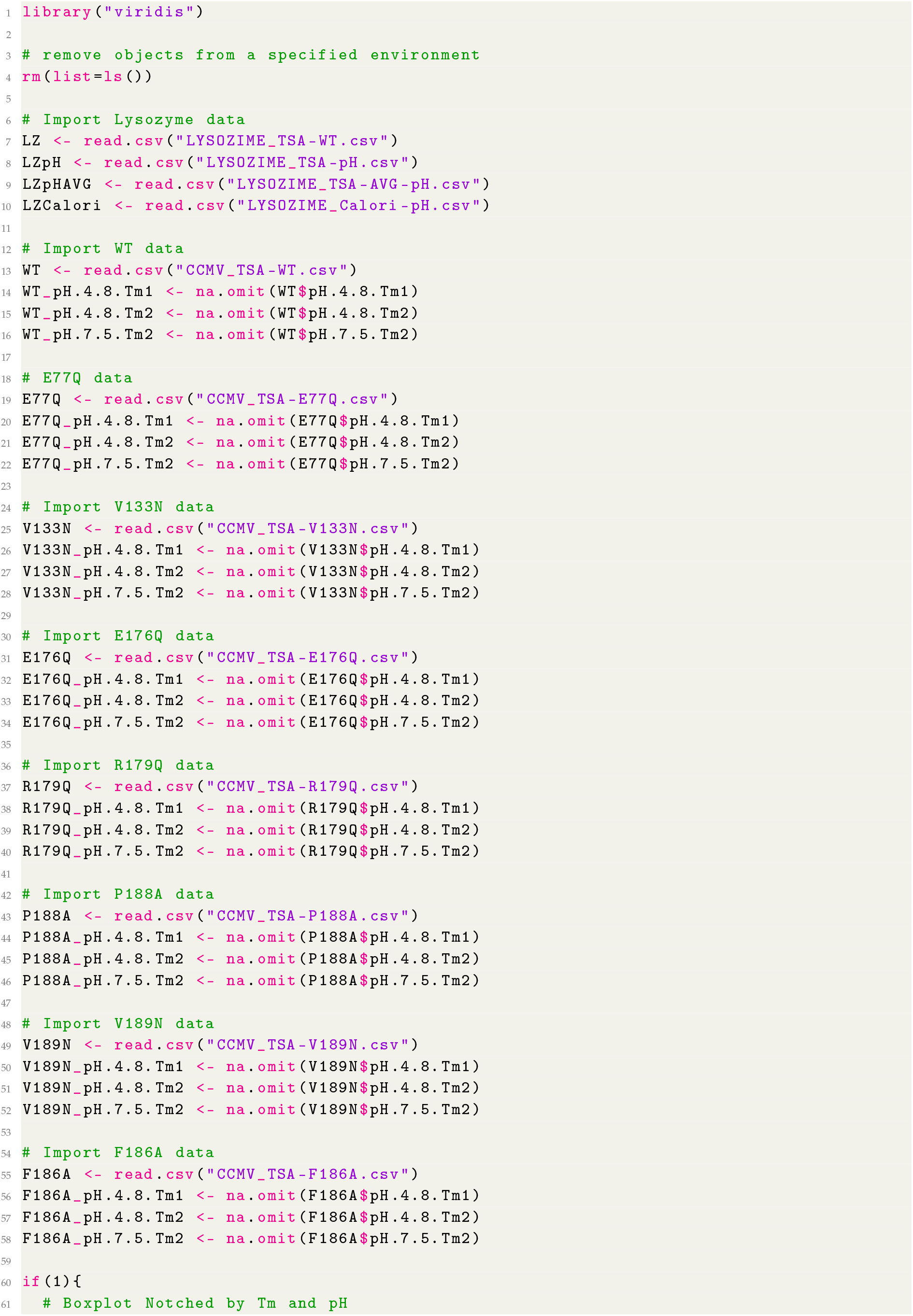

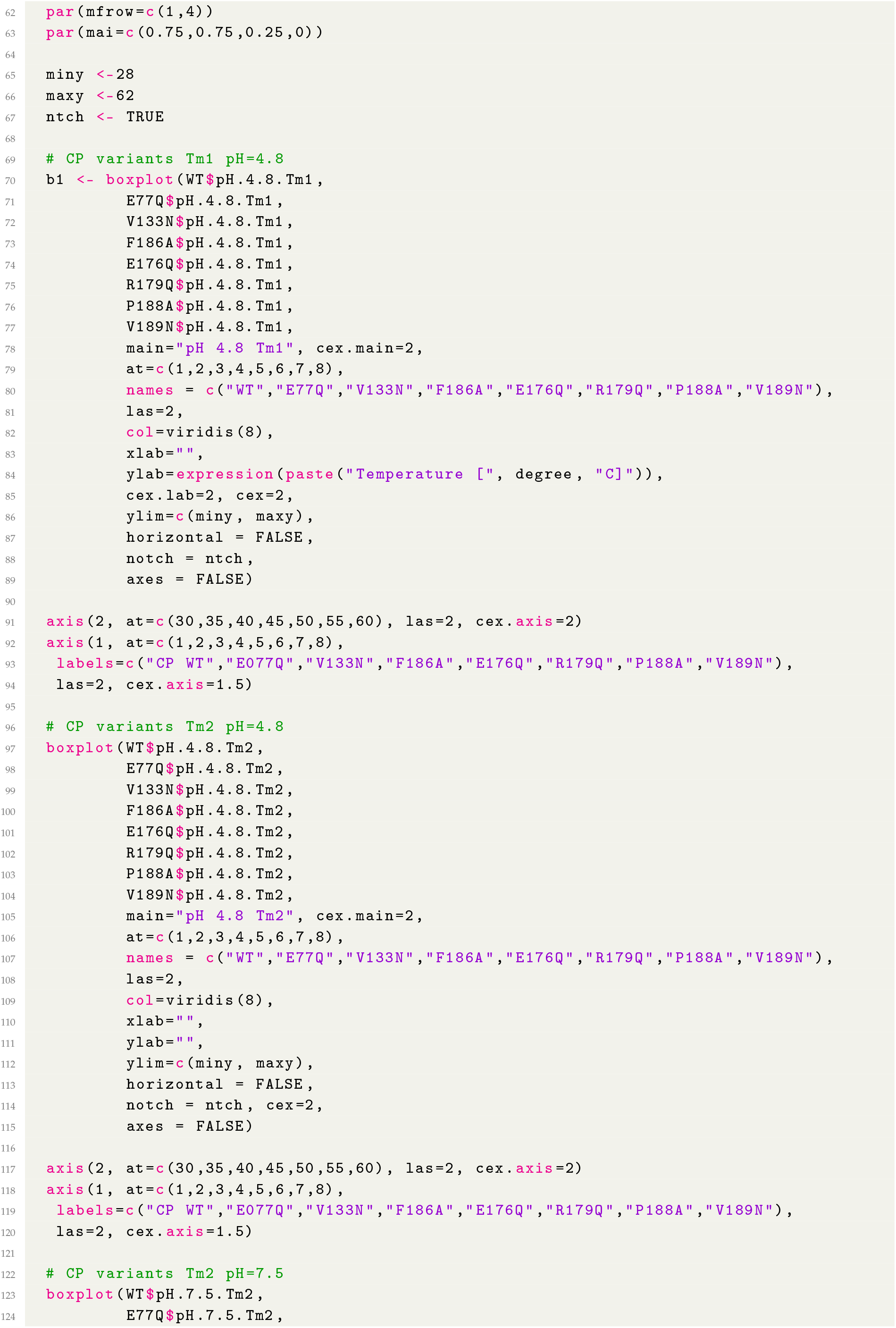

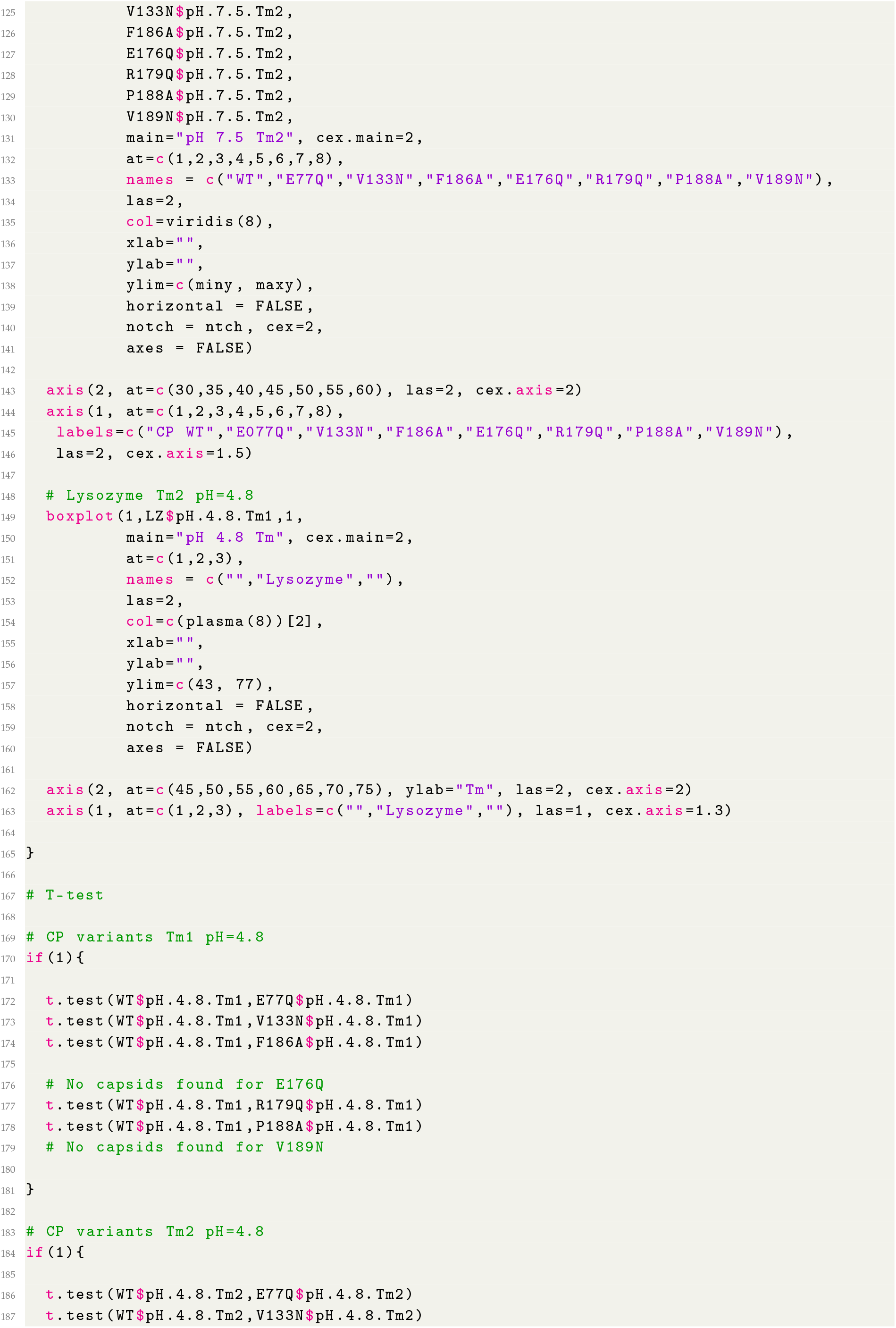

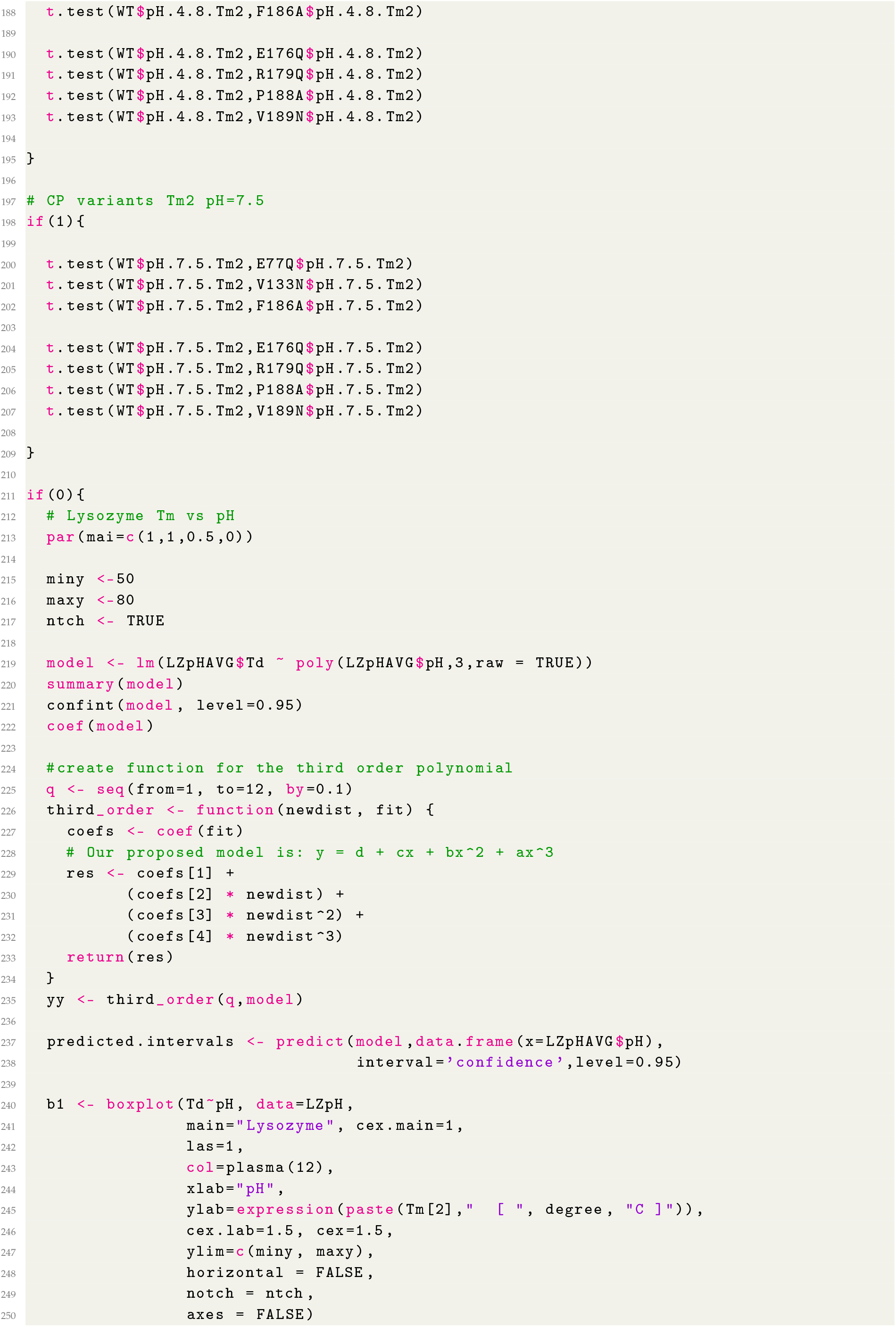

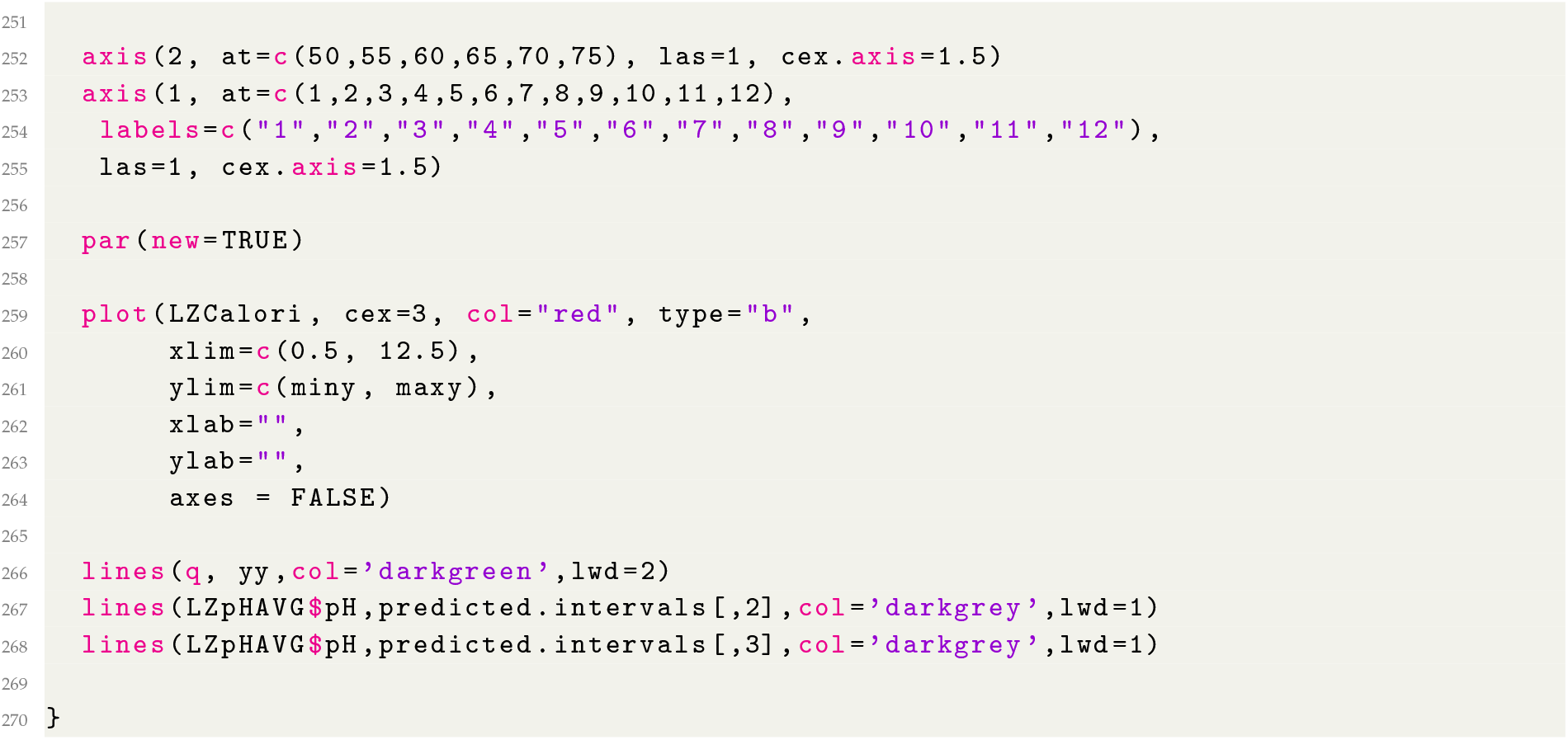

